# Effects of netarsudil-family Rho kinase inhibitors on human trabecular meshwork cell contractility and actin remodeling using a bioengineered ECM hydrogel

**DOI:** 10.1101/2022.05.19.492652

**Authors:** Tyler Bagué, Ayushi Singh, Rajanya Ghosh, Hannah Yoo, Curtis Kelly, Mitchell A. deLong, Casey C. Kopczynski, Samuel Herberg

**Author notes:** To whom correspondence should be addressed: Samuel Herberg, PhD, Assistant Professor; Department of Ophthalmology and Visual Sciences, SUNY Upstate Medical University, 505 Irving Avenue, Neuroscience Research Building Room 4609, Syracuse, NY 13210, USA.

## Abstract

Interactions between trabecular meshwork (TM) cells and their extracellular matrix (ECM) are critical for normal outflow function in the healthy eye. Multifactorial dysregulation of the TM is the principal cause of elevated intraocular pressure that is strongly associated with glaucomatous vision loss. Key characteristics of the diseased TM are pathologic contraction and actin stress fiber assembly, contributing to overall tissue stiffening. Among first-line glaucoma medications, the Rho-associated kinase inhibitor (ROCKi) netarsudil is known to directly target the stiffened TM to improve outflow function via tissue relaxation involving focal adhesion and actin stress fiber disassembly. Yet, no *in vitro* studies have explored the effect of netarsudil on human TM (HTM) cell contractility and actin remodeling in a 3D ECM environment. Here, we use our bioengineered HTM cell-encapsulated ECM hydrogel to investigate the efficacy of different netarsudil-family ROCKi compounds on reversing pathologic contraction and actin stress fibers. Netarsudil and all related experimental ROCKi compounds exhibited significant ROCK1/2 inhibitory and focal adhesion disruption activities. Furthermore, all ROCKi compounds displayed potent contraction-reversing effects on HTM hydrogels upon glaucomatous induction in a dose-dependent manner, relatively consistent with their biochemical/cellular inhibitory activities. At their tailored EC_50_ levels, netarsudil-family ROCKi compounds exhibited distinct effect signatures of reversing pathologic HTM hydrogel contraction and actin stress fibers, independent of the cell strain used. Netarsudil outperformed the experimental ROCKi compounds in support of its clinical status. In contrast, at uniform EC_50_-levels using netarsudil as reference, all ROCKi compounds performed similarly. Collectively, our data suggest that netarsudil exhibits high potency to rescue HTM cell pathobiology in a tissue-mimetic 3D ECM microenvironment, solidifying the utility of our bioengineered hydrogel model as a viable screening platform to further our understanding of TM pathophysiology in glaucoma.

## Introduction

The trabecular meshwork (TM) drains the aqueous humor to regulate outflow resistance, thereby maintaining normal intraocular pressure in the healthy eye (Overby et al., 2009; Tamm et al., 2015). The dynamic reciprocity between TM cells and their extracellular matrix (ECM) is critical in this process, as TM cells regulate tissue contraction and ECM remodeling to support outflow homeostasis (Kelley et al., 2009). In primary open-angle glaucoma, the most common form of glaucoma (Kwon et al., 2009), the TM undergoes increased fibrotic-like contraction, actin stress fiber assembly, ECM remodeling, and overall stiffening (Kwon et al., 2009; Wang et al., 2017). These cell-driven alterations lead to decreased trabecular outflow and consequently increased intraocular pressure, which if left untreated can push the TM to exceed its adaptive homeostatic capacity in a fed-forward loop (Stamer and Acott, 2012; Wang et al., 2017; Acott et al., 2021). The resulting TM dysfunction poses a serious threat to normal vision; approximately 80 million people worldwide are affected by glaucoma, a leading cause of blindness (Quigley, 1993; Quigley and Broman, 2006; Weinreb et al., 2014), and this number is projected to increase by almost 40% over the next 20 years (Tham et al., 2014).

Most ocular hypertension/glaucoma medications do not specifically target the diseased TM. They rather lower intraocular pressure by increasing uveoscleral outflow, bypassing the TM altogether, or by decreasing aqueous humor production (Beidoe and Mousa, 2012; Lin et al., 2018). In contrast, the FDA-approved Rho-associated kinase inhibitor (ROCKi) netarsudil, the active ingredient in Rhopressa™, increases outflow through the stiffened TM to increase outflow via reducing TM contraction as a function of ECM-focal adhesion and actin stress fiber disassembly (Rao et al., 2001; Zhang et al., 2012; Wang and Chang, 2014; Rao et al., 2017; Tanna and Johnson, 2018). The ability of netarsudil to improve outflow facility has been demonstrated in preclinical animal studies (Wang et al., 2015; Li et al., 2016), in an *ex vivo* perfusion study of human donor eyes (Ren et al., 2016), and in two human clinical studies (Kazemi et al., 2018; Sit et al., 2021). In contrast, only a single study to date has investigated netarsudil’s effect on normal and glaucomatous human TM (HTM) cell actin remodeling *in vitro* using live cell-imaging (Keller and Kopczynski, 2020). To our knowledge, no study has directly assessed the effect of netarsudil on HTM cell contractility in a relevant 3D ECM environment.

In the juxtacanalicular tissue region, TM cells reside embedded within a soft 3D ECM comprised of fibrillar and non-fibrillar collagens, elastic fibrils, glycosaminoglycans, proteoglycans, and matricellular proteins (Acott and Kelley, 2008; Tamm, 2009; Hann and Fautsch, 2011; Keller and Acott, 2013; Abu-Hassan et al., 2014). This is in stark contrast to conventional 2D tissue culture substrates that are known to create non-physiological culture conditions (Caliari and Burdick, 2016; Jensen and Teng, 2020). To that end, ECM biopolymer hydrogels provide a favorable tissue-mimetic 3D microenvironment and facilitate accurate *in vitro* modeling of cellular behaviors (Li et al., 2021). We recently reported a bioengineered ECM hydrogel composed of donor-derived HTM cells encapsulated within ECM biopolymers native to the TM to more accurately recapitulate the juxtacanalicular tissue region under normal and simulated glaucomatous conditions (Li et al., 2022a; Li et al., 2022b). Importantly, our model enables correlative analyses of TM cell cytoskeletal organization with tissue-level functional changes such as pathologic contraction contingent on 3D TM cell-ECM interactions.

Here, we investigate the effects of clinically-used netarsudil and different netarsudil-family experimental ROCKi compounds on reversing transforming growth factor beta 2 (TGFβ2)-induced (Inatani et al., 2001; Agarwal et al., 2015) pathologic HTM cell contractility and actin remodeling using our bioengineered ECM hydrogel.

## Materials and Methods

### HTM cell isolation and culture

Human donor eye tissue use was approved by the SUNY Upstate Medical University Institutional Review Board (protocol #1211036), and all experiments were performed in accordance with the tenets of the Declaration of Helsinki for the use of human tissue. Primary human TM (HTM) cells were isolated from healthy donor corneal rims discarded after transplant surgery, as previously described (Li et al., 2021; Li et al., 2022a; Li et al., 2022b), and cultured according to established protocols (Stamer et al., 1995; Keller et al., 2018). Three normal HTM cell strains (HTM05 [Male/57], HTM07 [Male/39], HTM36 [Female/56]) were used for the experiments in this study. All HTM cell strains were validated with dexamethasone-induced (DEX; Fisher Scientific, Waltham, MA, USA; 100 nM) myocilin expression in more than 50% of cells by immunocytochemistry and immunoblot analyses (**Suppl. Fig. 1**). All studies were conducted using cell passage 3-7. HTM cells were cultured in low-glucose Dulbecco’s Modified Eagle’s Medium (DMEM; Gibco; Thermo Fisher Scientific) containing 10% fetal bovine serum (FBS; Atlanta Biologicals, Flowery Branch, GA, USA) and 1% penicillin/streptomycin/glutamine (PSG; Gibco) and maintained at 37°C in a humidified atmosphere with 5% CO_2_. Fresh media was supplied every 2-3 days.

### Rho kinase inhibitors

Netarsudil (AR-13324), compound A (AR-13540), compound B (AR-16257), compound C (AR-13533), and compound D (AR-12862) were synthesized at Aerie Pharmaceuticals Inc., Durham, NC, USA (**Fig. 1**). All stocks were provided at 1.0 mM in dimethyl sulfoxide (DMSO). The solutions were sterilized using 0.2 µm syringe filters.

**Fig. 1.**
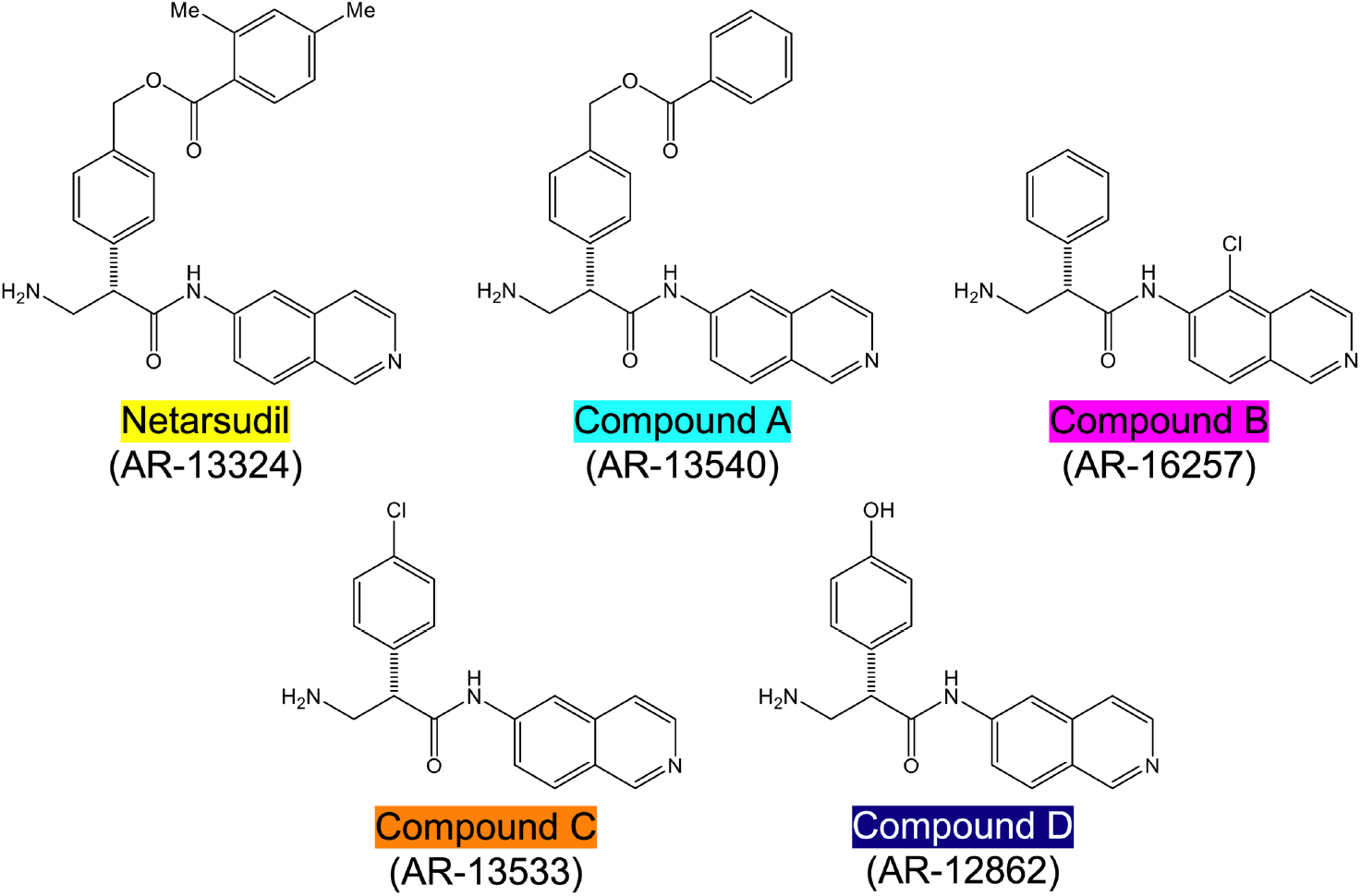
Structures of netarsudil-family ROCKi test compounds. Color code used throughout all figures.

### Biochemical and cell-based activity assays

Protein kinase assays (ROCK1 and ROCK2; from Invitrogen; Thermo Fisher Scientific) were conducted using serially diluted netarsudil/compounds A-D, as previously described (Sturdivant et al., 2016). In brief, ROCK1/2 activity was quantitated in 96-well white, flatbottom, half-area, nonbinding assay plates (No. 3642; Corning; Sigma-Aldrich) using the Kinase-Glo® Luminescent Kinase Assay (Promega, Madison, WI, USA) according to the manufacturer’s instructions. Dose response analyses were conducted to establish IC_50_ values, which were converted to K_i_ values using the Cheng-Prusoff Equation (K_i_ = IC_50_ / (1 + ([ATP]/Km ATP)).

Cell-based assays measuring disruption of focal adhesions were conducted using transformed HTM cells (TM-1; kind gift from Donna Peters; Department of Ophthalmology and Visual Sciences, University of Wisconsin), as previously described (Sturdivant et al., 2016). In brief, HTM cells were grown on fibronectin-coated glass-bottom 96-well plates and incubated in media containing serially diluted netarsudil/compounds A-D for 6 h, followed by fixing in formaldehyde and routine processing for immunocytochemistry. HTM cells were stained with an anti-paxillin primary antibody followed by incubation with an Alexa Fluor-488 fluorescent secondary antibody and Hoechst 33342 counterstain (all from Invitrogen) to reveal focal adhesions and nuclei, respectively. Images were collected on an INCell 1000 imager (GE Healthcare, Marlborough, MA, USA), and total area of focal adhesions was measured using a custom algorithm developed using the INCell Developer Toolbox, v1.6. Dose response analyses were conducted to establish IC_50_ values.

### Hydrogel precursor solutions

Methacrylate-conjugated bovine collagen type I (MA-COL; Advanced BioMatrix, Carlsbad, CA, USA) was reconstituted in sterile 20 mM acetic acid at 6 mg/ml. Immediately prior to use, 1 ml MA-COL was neutralized with 85 µl neutralization buffer (Advanced BioMatrix) according to the manufacturer’s instructions. Thiol-conjugated hyaluronic acid (SH-HA; Glycosil®; Advanced BioMatrix) was reconstituted in sterile diH_2_O containing 0.5% (w/v) photoinitiator (4-(2-hydroxyethoxy) phenyl-(2-propyl) ketone; Irgacure® 2959; Sigma-Aldrich, St. Louis, MO, USA) at 10 mg/ml according to the manufacturer’s protocol. In-house expressed elastin-like polypeptide (SH-ELP; thiol via cysteine in K<u>C</u>TS flanks (Zhang et al., 2015; Li et al., 2021)) was reconstituted in DPBS at 10 mg/ml and sterilized using a 0.2 µm syringe filter in the cold. The photoactive ECM biopolymers can form chemical crosslinks via methacrylate, thiol-ester, or disulfide linkages.

### Preparation of HTM hydrogels

HTM cells (1.0 × 10^6^ cells/ml) were thoroughly mixed with MA-COL (3.6 mg/ml), SH-HA (0.5 mg/ml, 0.025% (w/v) photoinitiator) and SH-ELP (2.5 mg/ml) on ice (**Suppl. Fig. 2A**), followed by pipetting 10 μl droplets of the HTM cell-laden hydrogel precursor solution onto polydimethylsiloxane-coated (PDMS; Sylgard 184; Dow Corning) 24-well culture plates (**Suppl. Fig. 2B**), according to our established protocols (Li et al., 2021; Li et al., 2022a; Li et al., 2022b). Alternatively, 30 µl droplets of the HTM cell-laden hydrogel precursor solution were pipetted onto Surfasil-coated (Fisher Scientific) 18 × 18-mm square glass coverslips followed by placing a regular 12-mm round glass coverslip onto the hydrogels to facilitate even spreading of the polymer solution. HTM hydrogels were crosslinked by exposure to UV light (OmniCure S1500 UV Spot Curing System; Excelitas Technologies, Mississauga, Ontario, Canada) at 320-500 nm, 2.2 W/cm^2^ for 5 s. The HTM hydrogel-adhered coverslips were removed with fine-tipped tweezers and placed hydrogel-side facing up in PDMS-coated 24-well culture plates (**Suppl. Fig. 2C**).

### HTM hydrogel treatments

HTM hydrogels were cultured in DMEM with 10% FBS and 1% PSG, and subjected to the following treatments for 10 d: **1) Control** (vehicle: 40 μM HCL, 0.002% BSA [0-5 d]; 0.1% DMSO [5-10 d]; all from Thermo Fisher Scientific) for 0-10 d <u>[=Baseline]</u>, **2) TGFβ2** (TGFβ2: 2.5 ng/ml; R&D Systems, Minneapolis, MN, USA) for 0-5 d followed by vehicle control (0.1% DMSO) for 5-10 d <u>[=Induction]</u>, and **3) TGFβ2 + ROCKi** (TGFβ2: 2.5 ng/ml; R&D Systems) for 0-5 d followed by netarsudil/compounds A-D (0.0001 µM, 0.001 µM, 0.01 µM, 0.1 µM, 1.0 µM; Aerie Pharmaceuticals) or Y27632 (10 μM; Sigma-Aldrich) <u>[=Induction + Rescue]</u> (**Fig. 2**). First, dose response analyses (i.e., the ability to reduce TGFβ2-induced HTM hydrogel contraction) were conducted with netarsudil and compounds A-D to establish EC_50_ values using one HTM cell strain (i.e., HTM07); Y27632 served as a reference control. Subsequently, netarsudil was directly compared with compounds A-D at their respective EC_50_ using three HTM cell strains (i.e., HTM05, HTM07, HTM36), with netarsudil at 1.0 µM serving as a reference control.

**Fig. 2.**
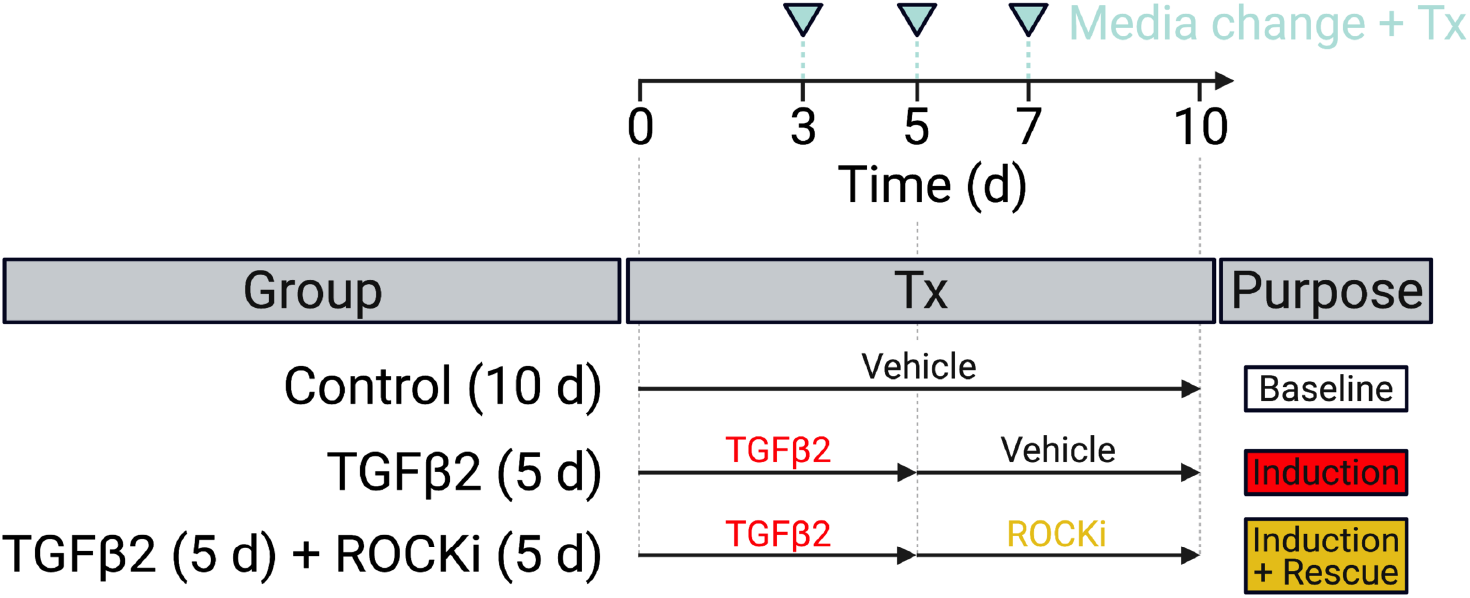
Experimental design. HTM hydrogels were treated with vehicle control (=baseline) or TGFβ2 for 5 d to induce a glaucoma-like cell phenotype (=induction) before adding netarsudil-family ROCKi test compounds for the next 5 d in absence of TGFβ2 (=induction + rescue), with fresh media supplied every 2-3 days.

### HTM hydrogel contraction analysis

Longitudinal brightfield images of HTM hydrogels subjected to the different treatments were acquired at 0 d and 10 d with an Eclipse T*i* microscope (Nikon Instruments, Melville, NY, USA). Construct area was measured using Fiji software (National Institutes of Health (NIH), Bethesda, MD, USA) and normalized to 0 d followed by normalization to controls.

### HTM hydrogel cell viability analysis

The number of viable cells inside HTM hydrogels subjected to the different treatments for 10 d was measured with the CellTiter 96® Aqueous Non-Radioactive Cell Proliferation Assay (MTS; Promega) according to the manufacturer’s instructions. HTM hydrogels were incubated with the staining solution (38 μl MTS, 2 μl PMS solution, 200 μl DMEM) at 37ºC for 1.5 h. Absorbance at 490 nm was recorded using a spectrophotometer plate reader (BioTek, Winooski, VT, USA). Blank (DMEM with the staining solution)-subtracted absorbance values served as a direct measure of HTM cell viability.

### HTM hydrogel immunocytochemistry analysis

HTM hydrogels on coverslips subjected to the different treatments for 10 d were fixed with 4% paraformaldehyde (Thermo Fisher Scientific) at 4ºC overnight, permeabilized with 0.5% Triton™ X-100 (Thermo Fisher Scientific), blocked with blocking buffer (BioGeneX), and incubated with Phalloidin-iFluor 488 to stain for F-actin (Abcam, Cambridge, MA, USA) and a Cy3-conjugated primary antibody against α-smooth muscle actin (anti-αSMA [C6198] 1:500; Sigma-Aldrich). Nuclei were counterstained with 4′,6′-diamidino-2-phenylindole (DAPI; Abcam). Coverslips were mounted with ProLong™ Gold Antifade (Invitrogen) on Superfrost™ microscope slides (Fisher Scientific), and fluorescent images were acquired with a Zeiss LSM 780 confocal microscope (Zeiss, Germany). The image size was set to 512 × 512 pixels in x/y with a resolution of 1.66 μm per pixel. Individual z-stacks consisted of 7 slices with the z-step interval set to 13.3 μm. Fluorescence signal intensity of F-actin and αSMA were determined using Z-project maximum intensity projections in FIJI (NIH) with image background subtraction.

### Statistical analysis

Individual sample sizes are specified in each figure caption. Comparisons between groups were assessed by one-way or two-way analysis of variance (ANOVA) with Tukey’s multiple comparisons *post hoc* tests, as appropriate. All data are shown with mean ± SD, some with individual data points. The significance level was set at p<0.05 or lower. GraphPad Prism software v9.3 (GraphPad Software, La Jolla, CA, USA) was used for all analyses.

## Results

### In vitro activity of netarsudil-family ROCK inhibitors

First, we investigated the specific inhibitory activity of different netarsudil-family ROCKi against the two human Rho kinase isoforms ROCK1 and ROCK2, as well as the compounds’ ability to disrupt focal adhesions in HTM cells according to established protocols (Lin et al., 2018). All ROCKi compounds exhibited significant inhibitory activity against ROCK1 and ROCK2, with the order of potency from highest to lowest as follows: compound A>compound C>compound B>netarsudil>compound D (**Table 1**). The rank order of potency to disrupt HTM cell focal adhesions on conventional 2D culture substrates was: compound C>compound A>netarsudil>compound D (**Table 1**).

**Table 1.** *In vitro* potency of netarsudil-family ROCK inhibitors.

| Compound | ROCK1 K<sub>i</sub> (nM) | ROCK2 K<sub>i</sub> (nM) | HTM IC<sub>50</sub> (nM) |
| --- | --- | --- | --- |
| Netarsudil | 2.6 ± 0.7 | 2.8 ± 0.7 | 17 ± 2 |
| Compound A | 1.5 ± 0.7 | 1.2 ± 0.3 | 15 ± 3 |
| Compound B | 1.7 ± 0.0* | 2.0 ± 0.0* | Not tested |
| Compound C | 1.8 ± 0.3 | 1.2 ± 0.4 | 5.6 ± 0.7 |
| Compound D | 4.5 ± 2.0 | 1.7 ± 0.7 | 193 ± 51 |
| Mean ± SEM, N ≥ 3, *N = 2; K<sub>i</sub>, inhibition constant; IC<sub>50</sub>, half maximal inhibitory concentration | | | |

Together, these data show that netarsudil-family ROCKi treatments exhibit potent ROCK1/2 inhibitory and focal adhesion disruption activities. The differential responses observed with clinically-used netarsudil compared to experimental compounds A-D likely stems from differences in compound structure/chemistry (**Fig. 1**), affecting cellular uptake and efficacy.

### Dose response analysis of netarsudil-family ROCK inhibitors on reversing TGFβ2-induced HTM hydrogel contraction

The contractility status of the TM influences outflow resistance and intraocular pressure (Dismuke et al., 2014). In our recent study, we showed that HTM cells encapsulated in ECM biopolymer hydrogels are highly contractile, and that TGFβ2 increases HTM hydrogel contraction compared to vehicle-treated controls (Li et al., 2021).

Therefore, to investigate the potency of different netarsudil-family ROCKi compounds in rescuing pathologic HTM hydrogel contraction, constructs were treated with TGFβ2 for 5 d to induce a glaucoma-like cell phenotype before adding netarsudil/compounds A-D over a broad dose range for the next 5 d in the absence of TGFβ2. Treatment with TGFβ2 significantly increased HTM hydrogel contraction compared to controls (=lower values; **Fig. 3A-J**), consistent with our previous reports (Li et al., 2021; Li et al., 2022a; Li et al., 2022b). All ROCKi treatments reversed TGFβ2-induced contraction in a dose-dependent manner (=higher values). For netarsudil, we observed significantly decreased HTM hydrogel contraction using 0.01 µM and higher concentrations compared to the TGFβ2 group in a near-linear fashion. Netarsudil at 0.1 µM was equivalent to the standard concentration of 10 µM Y27632 (i.e., 100x more concentrated), whereas 1.0 µM netarsudil was significantly more potent compared to Y27632 (**Fig. 3A,B**). The EC_50_ for netarsudil was 35.9 nM (**Fig. 3K**). Similarly, for compound A, we found significantly decreased TGFβ2-induced HTM hydrogel contraction at 0.01 µM and above, plateauing at 0.1 µM. Consequently, even at 1.0 µM compound A was not different from 10 µM Y27632 (**Fig. 3C,D**). The EC_50_ for compound A was 3.7 nM (**Fig. 3K**). For compound B, we observed significantly decreased HTM hydrogel contraction only at 0.1 µM and 1.0 µM compared to the TGFβ2 group, with behavior at 1.0 µM being comparable to compound A (**Fig. 3E,F**). The EC_50_ for compound B was 22.2 nM (**Fig. 3K**). For compound C, we found significantly decreased TGFβ2-induced HTM hydrogel contraction using 0.01 µM and above, with a noticeable “jump” between 0.001 µM and 0.01 µM. Of note, even at 1.0 µM compound C was comparable to standard 10 µM Y27632 (**Fig. 3G,H**), which showed some variability between experiments. The EC_50_ for compound C was 6.7 nM (**Fig. 3K**). Lastly, for compound D, we observed significantly decreased HTM hydrogel contraction only at 0.1 µM and 1.0 µM compared to the TGFβ2 group – similar to compound B – with a more “blunted” response over the tested dose range compared to all other ROCKi treatments (**Fig. 3I,J**). Consequently, the EC_50_ for compound D was 123.3 nM (**Fig. 3K**).

**Fig. 3.**
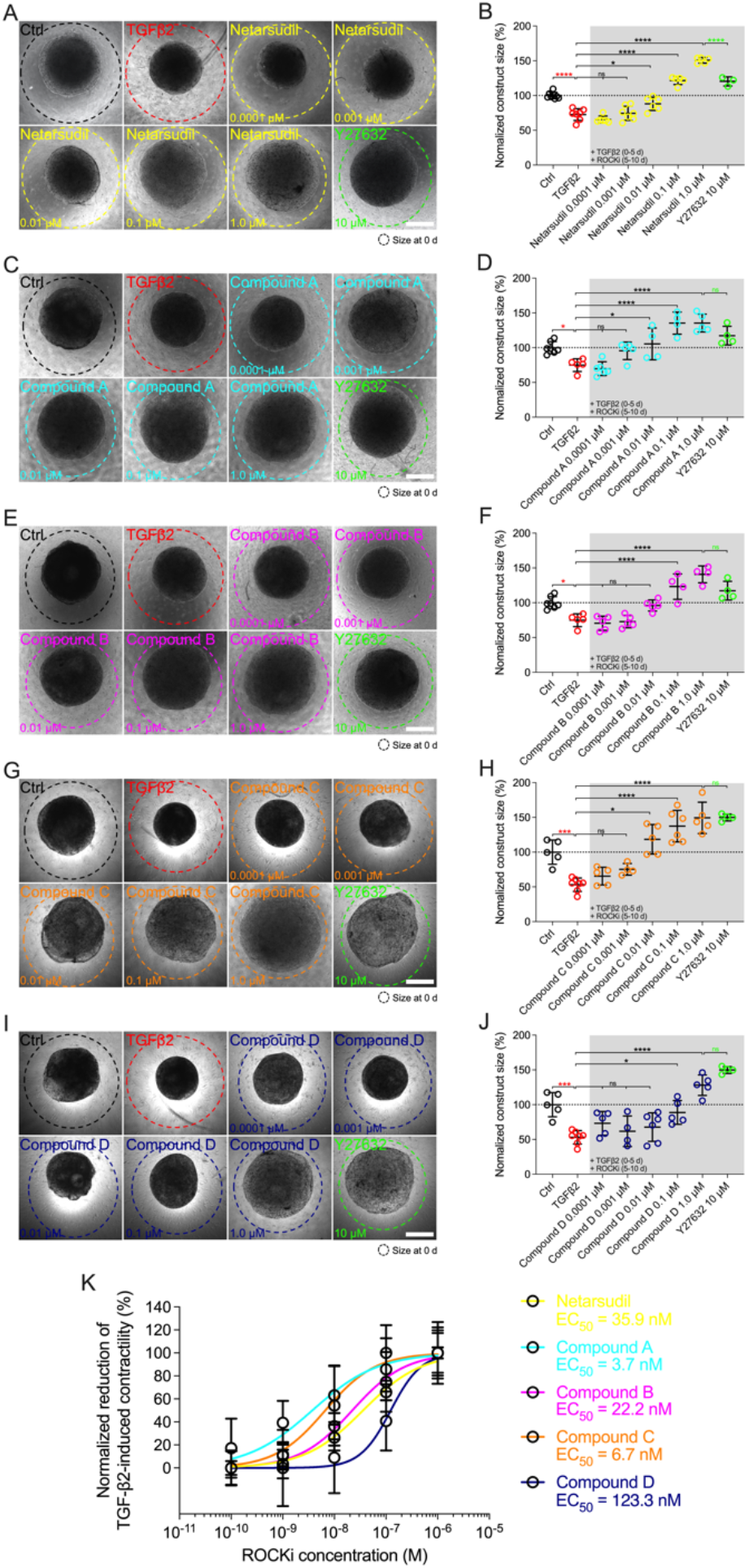
Dose response effects of netarsudil-family ROCKi treatment following TGFβ2-induction on HTM hydrogel contraction. Representative brightfield micrographs of HTM hydrogels encapsulated with HTM07 subjected to (**A**) netarsudil, (**C**) compound A, (**E**) compound B, (**G**) compound C, or (**I**) compound D over a broad dose range for 10 d, with Y27632 serving as refence control (dashed lines outline original construct size at 0 d). Scale bars, 1 mm. Construct size quantification of HTM hydrogels subjected to (**B**) netarsudil, (**D**) compound A, (**F**) compound B, (**H**) compound C, or (**J**) compound D (N = 3-8 replicates per group). (**K**) ROCKi dose response curves with calculated EC_50_ values. Data shown as Mean ± SD with individual data points. Significance was determined by one-way ANOVA using multiple comparisons tests (*p<0.05, ***p<0.001,****p<0.0001; ns = not significant).

Together, these data show that netarsudil-family ROCKi treatments exhibit potent contraction-reversing effects on HTM cell-encapsulated ECM hydrogels upon glaucomatous induction in a dose-dependent manner. The order of potency from highest to lowest was as follows: compound A>compound C>compound B>netarsudil>compound D, precisely matching the compounds’ independently acquired ROCK1/2 inhibitory activities (**Table 1**).

### Comparison of netarsudil-family ROCK inhibitors at EC_50_ on reversing TGFβ2-induced HTM hydrogel contraction

Next, we directly compared the different netarsudil-family ROCKi compounds at their respective EC_50_ to assess the tailored efficacy in rescuing TGFβ2-induced HTM hydrogel contraction using HTM05 (**Fig. 4A,B**), HTM07 (**Fig. 4C,D**), and HTM36 (**Fig. 4E,F**). Of note, HTM05 and HTM36 cells showed an overall greater inherent contractility (i.e., vehicle-treated controls) compared to HTM07 (**Fig. 4A,C,E**), consistent with normal donor-to-donor variability observed in previous studies (Li et al., 2021). Treatment with TGFβ2 significantly increased HTM hydrogel contraction by ~25% over controls, independent of the cell strain used (**Fig. 4A-G**). Netarsudil at 35.9 nM significantly decreased TGFβ2-induced HTM hydrogel contraction, reaching ~120% across all three cell strains. While this overall effect was significantly less compared to max level netarsudil at 1.0 µM (=reference control and highest values; **Fig. 4G**), the difference in efficacy (~1.2-fold) was not as large as the order-of-magnitude lower dose would suggest. Compound A at 3.7 nM failed to reverse HTM hydrogel contraction compared to the TGFβ2 group in all HTM cell strains tested (**Fig. 4A-G**). Compound B at 22.2 nM significantly decreased TGFβ2-induced HTM hydrogel contraction using HTM05 (**Fig. 4A,B**) and HTM36 cells (**Fig. 4E,F**), reaching baseline levels; yet, the overall rescuing effect was less compared to netarsudil-EC_50_ (**Fig. 4G**). Compound C at 6.7 nM failed to rescue HTM hydrogel contraction compared to the TGFβ2 group across HTM cell strains (**Fig. 4A-G**), showing similar behavior as compound A. Lastly, compound D at 123.3 nM significantly decreased TGFβ2-induced HTM hydrogel contraction using HTM05 (**Fig. 4A,B**) and HTM36 cells (**Fig. 4E,F**), approximating baseline levels. The overall rescuing effect was equivalent to compound B but less compared to netarsudil-EC_50_ (**Fig. 4G**). To rule out that HTM hydrogel contractility was influenced by the cell number, we assessed HTM cell viability in constructs subjected to the different treatments at the experimental end point. No differences were noted across treatment groups for all HTM cell strains (**Suppl. Fig. 3**).

**Fig. 4.**
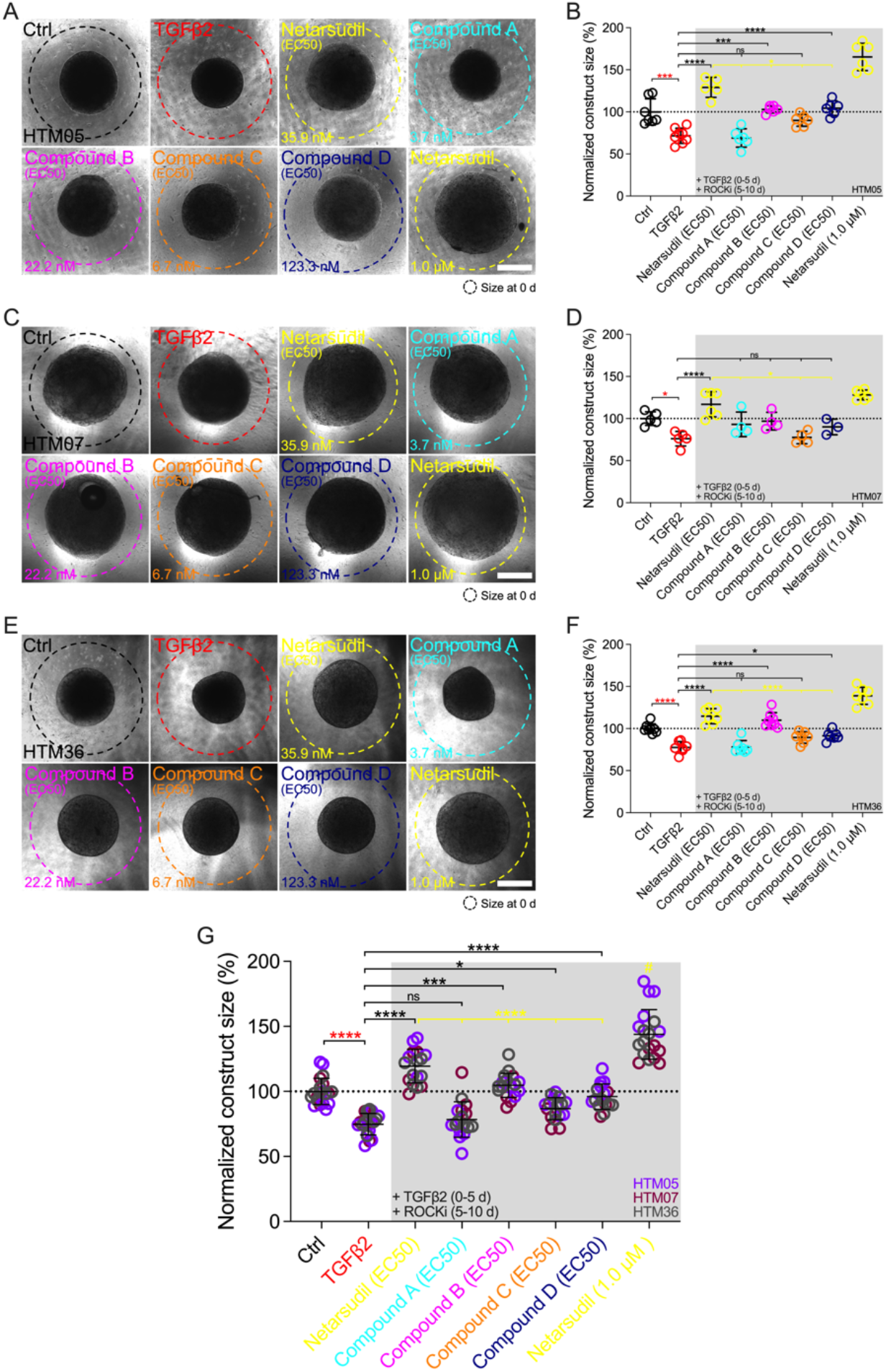
Effects of netarsudil-family ROCKi treatment at EC_50_ following TGFβ2-induction on HTM hydrogel contraction. Representative brightfield micrographs of HTM hydrogels encapsulated with (**A**) HTM05, (**C**) HTM07, or (**E**) HTM36 subjected to the different treatments for 10 d, with max level netarsudil serving as refence control (dashed lines outline original construct size at 0 d). Scale bars, 1 mm. Construct size quantification of HTM hydrogels encapsulated with (**B**) HTM05, (**D**) HTM07, or (**F**) HTM36 (N = 3-8 replicates per group). (**G**) Construct size quantification of HTM hydrogels encapsulated with HTM05 (purple), HTM07 (maroon), or HTM36 (gray). Data shown as Mean ± SD with individual data points. Significance was determined by one- or two-way ANOVA using multiple comparisons tests (*p<0.05, ***p<0.001, ****p<0.0001; ns = not significant; ^#^p<0.0001 vs. all other groups).

Together, these data show that the different netarsudil-family ROCKi treatments at <u>tailored</u> EC_50_-levels exhibit distinct signatures of rescuing pathologic HTM hydrogel contraction, independent of the HTM cell strain used. The overall order of efficacy from highest to lowest was as follows: netarsudil>compound B>compound D>compound C>compound A. Netarsudil at both EC_50_ and max levels outperformed the experimental compounds A-D, thereby validating its clinical status in our soft tissue-mimetic 3D ECM hydrogel system.

### Comparison of netarsudil-family ROCK inhibitors at EC_50_ on reversing TGFβ2-induced HTM cell actin cytoskeletal remodeling

The actin cytoskeleton is the primary force-generating machinery, with filamentous (F)-actin fiber arrangement directly affecting cell and tissue contraction (Svitkina, 2018). F-actin and alpha smooth muscle actin (αSMA) fibers are involved in HTM cell contractility regulation, and we have previously demonstrated that their abundance and organization is altered in hydrogel-encapsulated HTM cells under simulated glaucomatous conditions (Li et al., 2021).

Therefore, we next investigated the efficacy of different netarsudil-family ROCKi compounds at their respective EC_50_ to reverse TGFβ2-induced HTM cell actin stress fiber formation within 3D ECM hydrogels using the same three different cell strains. Treatment with TGFβ2 significantly increased F-actin stress fiber formation/signal intensity compared to controls, independent of the cell strain used (**Fig. 5A-G**). Netarsudil at 35.9 nM significantly decreased TGFβ2-induced F-actin stress fibers using HTM05 (**Fig. 5A,B**) and HTM36 cells (**Fig. 5E,F**), restoring baseline levels (**Fig. 5G**). Similar to the contraction data, this overall effect was significantly less compared to netarsudil reference control at 1.0 µM (=lowest values). Compound A at 3.7 nM failed to reverse F-actin stress fibers compared to the TGFβ2 group using HTM05 (**Fig. 5A,B**) and HTM07 cells (**Fig. 5C,D**). While a significant reduction was noted using HTM36 (**Fig. 5E,F**), the overall response across cell strains was not different from TGFβ2 (**Fig. 5G**). Compound B at 22.2 nM significantly decreased TGFβ2-induced F-actin stress fibers using HTM05 (**Fig. 5A,B**) and HTM36 cells (**Fig. 5E,F**), reaching baseline levels, with the overall rescuing effect comparable to netarsudil-EC_50_ (**Fig. 5G**). Compound C at 6.7 nM failed to rescue F-actin stress fibers compared to the TGFβ2 group using HTM05 (**Fig. 5A,B**) and HTM07 cells (**Fig. 5C,D**). A significant reduction was noted using HTM36 (**Fig. 5E,F**), driving the overall response across cell strains to be significantly different from TGFβ2 and equivalent to netarsudil-EC_50_ (**Fig. 5G**). Lastly, compound D at 123.3 nM significantly decreased TGFβ2-induced F-actin stress fibers across all three HTM cell strains, with the overall rescuing effect being comparable to netarsudil-EC_50_ (**Fig. 5A-G**). Of note, while TGFβ2-treated HTM hydrogels showed qualitatively increased αSMA stress fiber formation/signal intensity compared to controls, no differences were noted across treatment groups for all HTM cell strains (**Suppl. Fig. 4**).

**Fig. 5.**
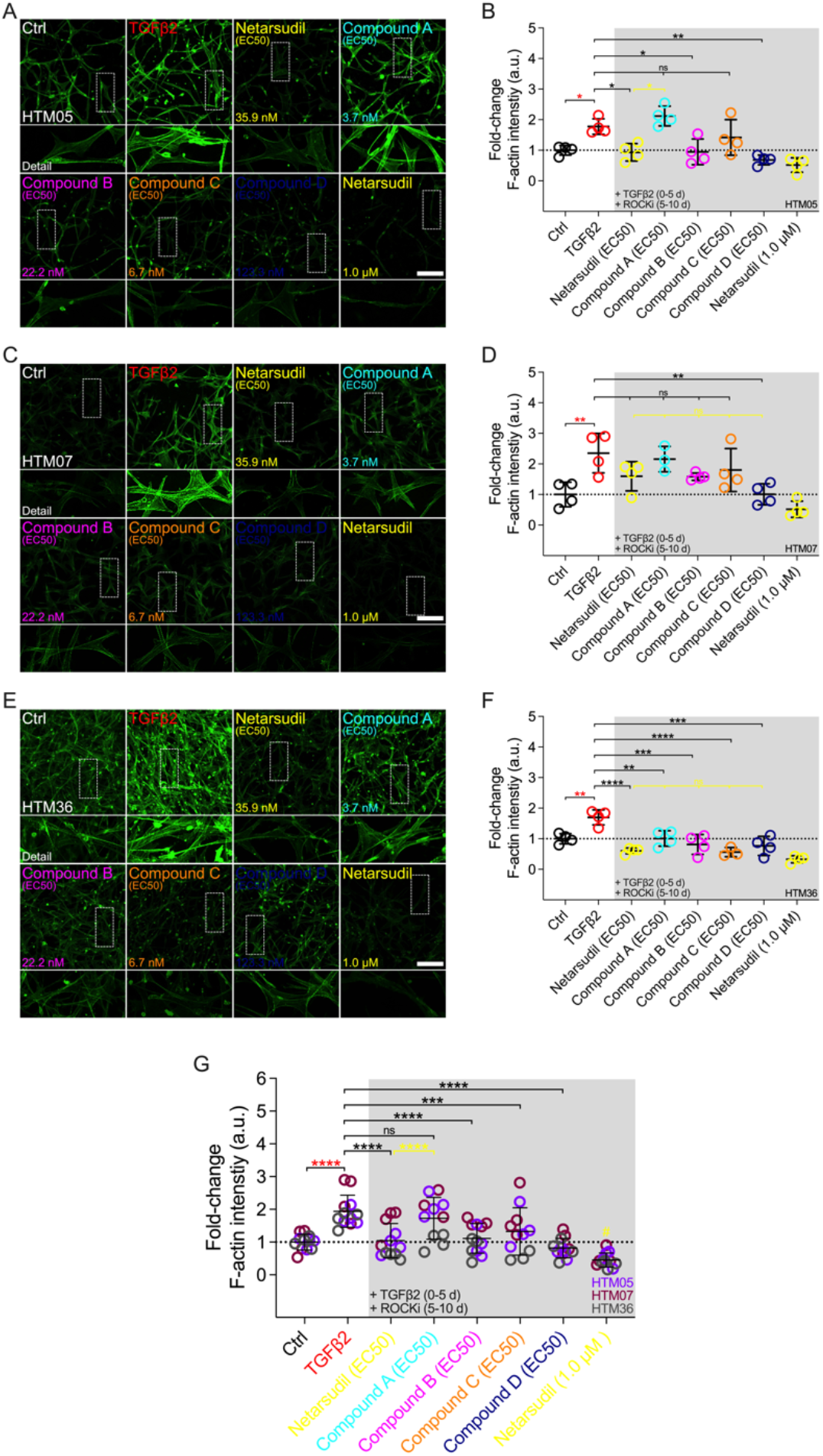
Effects of netarsudil-family ROCKi treatment at EC_50_ following TGFβ2-induction on HTM cell F-actin stress fibers within ECM hydrogels. Representative confocal fluorescence micrographs of F-actin in HTM hydrogels encapsulated with (**A**) HTM05, (**C**) HTM07, or (**E**) HTM36 subjected to the different treatments for 10 d (F-actin = green). Scale bar, 200 μm. Quantification of relative F-actin signal intensity in HTM hydrogels encapsulated with (**B**) HTM05, (**D**) HTM07, or (**F**) HTM36 (N = 4 replicates per group). (**G**) Pooled quantification of relative F-actin signal intensity in HTM hydrogels encapsulated with HTM05 (purple), HTM07 (maroon), or HTM36 (gray) (N = 4 replicates per group and HTM cell strain). Data shown as Mean ± SD with individual data points. Significance was determined by one- or two-way ANOVA using multiple comparisons tests (*p<0.05, **p<0.01, ***p<0.001,****p<0.0001; ns = not significant; ^#^p<0.01 vs. Ctrl, TGFβ2, netarsudil, and compounds A-C).

Together, these data show that irrespective of the HTM cell strain used the different netarsudil-family ROCKi treatments at <u>tailored</u> EC_50_-levels exhibit distinct effects of relaxing pathologic F-actin stress fibers in ECM hydrogel-encapsulated cells. The overall order of efficacy from highest to lowest was as follows: compound D>netarsudil>compound B>compound C>compound A. Netarsudil performed similarly to compounds B-D, with max level netarsudil nearly abolishing the F-actin signal.

### Comparison of netarsudil-family ROCK inhibitors at equivalent netarsudil-EC_50_ on reversing TGFβ2-induced HTM hydrogel contraction

To remove the potential confounder of variable ROCKi dosing, we compared the efficacy of the different netarsudil-family compounds in rescuing TGFβ2-induced HTM hydrogel contraction at a <u>uniform ROCKi concentration of 35.9 nM, the determined netarsudil-EC_50_</u>, using HTM07 (**Fig. 6A,B**) and HTM36 cells (**Fig. 6C,D**). Treatment with TGFβ2 significantly increased HTM hydrogel contraction by ~15% over controls for both cell strains (**Fig. 6A-E**). This was slightly less compared to earlier experiments, likely due to the necessity of using higher passage cells in this experiment; this trend of overall reduced hydrogel contractility continued across all treatment groups. As expected, netarsudil significantly decreased TGFβ2-induced HTM hydrogel contraction, restoring baseline in both cell strains (**Fig. 6A-E**). For HTM07, compounds A-D significantly decreased HTM hydrogel contraction compared to the TGFβ2 group reaching levels equivalent to netarsudil (**Fig. 6A,B,E**). For HTM36, compound A at a ~10-fold higher dose than earlier experiments significantly decreased TGFβ2-induced HTM hydrogel contraction, whereas compounds B-D failed to rescue pathologic contraction (**Fig. 6C,D**). The overall response across both HTM cell strains was that all ROCKi compounds significantly reversed TGFβ2-induced HTM hydrogel contraction, with no differences noted between netarsudil and compounds A-D (**Fig. 6E**).

**Fig. 6.**
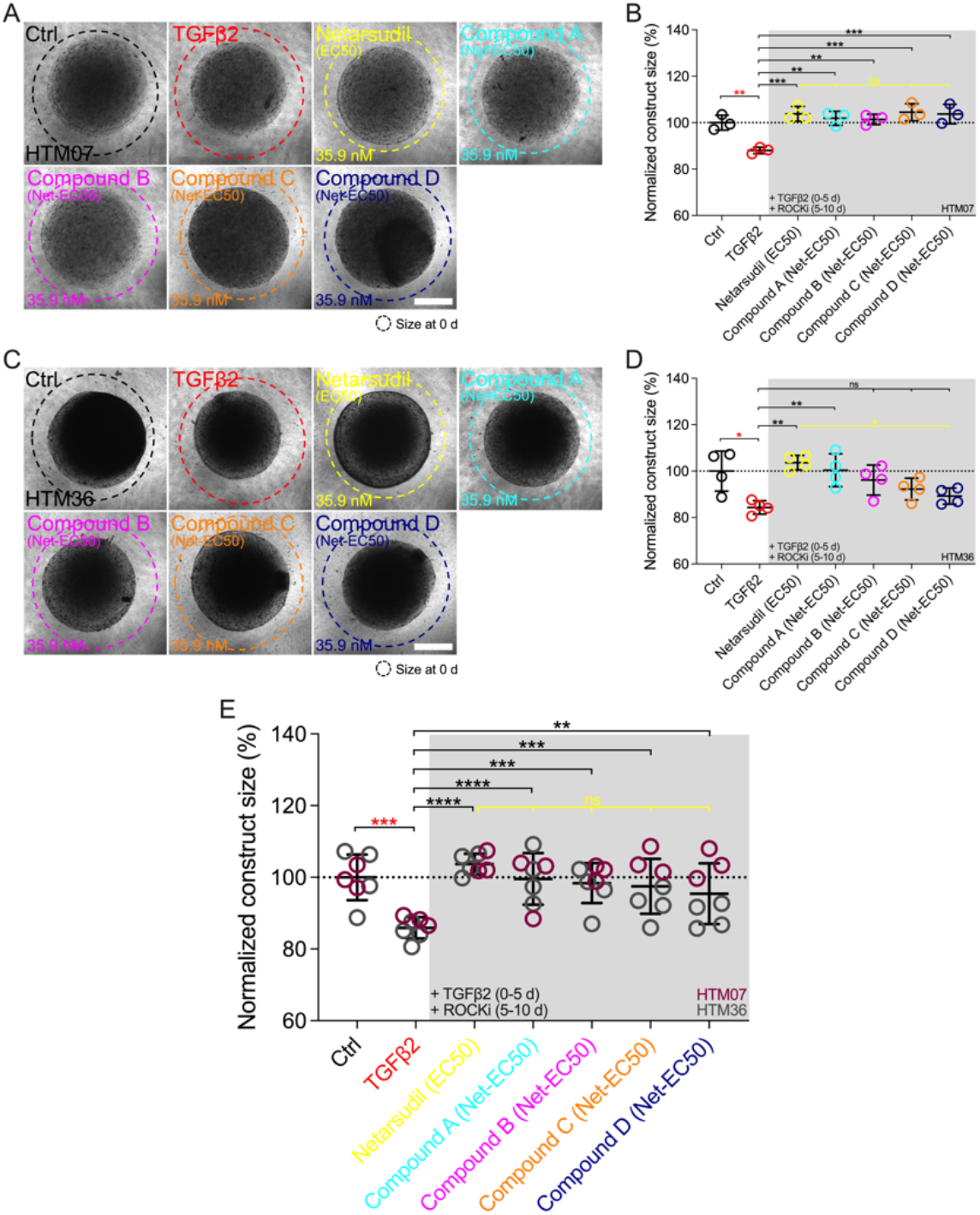
Effects of netarsudil-family ROCKi treatment at uniform netarsudil-EC_50_ following TGFβ2-induction on HTM hydrogel contraction. Representative brightfield micrographs of HTM hydrogels encapsulated with (**A**) HTM07 or (**C**) HTM36 subjected to the different treatments for 10 d (dashed lines outline original construct size at 0 d). Scale bars, 1 mm. Construct size quantification of HTM hydrogels encapsulated with (**B**) HTM07 or (**D**) HTM36 (N = 3-4 replicates per group). (**E**) Construct size quantification of HTM hydrogels encapsulated with HTM07 (maroon) or HTM36 (gray). Data shown as Mean ± SD with individual data points. Significance was determined by one- or two-way ANOVA using multiple comparisons tests (*p<0.05, **p<0.01, ***p<0.001, ****p<0.0001; ns = not significant).

Together, these data show that the different netarsudil-family ROCKi treatments at <u>uniform</u> EC_50_-levels lose their distinct signatures of rescuing pathologic HTM hydrogel contraction independent of the HTM cell strain used. When tested at equimolar dosing, netarsudil performed similarly to the compounds A-D, suggesting that subtle differences in experimental compound-specific activities can be offset by matching their doses with the clinical drug.

## Discussion

*In vitro* studies of TM cell function are fundamentally hampered by conventional cell culture systems. It is widely accepted that cells in 3D environments made of relevant ECM proteins frequently show divergent behavior compared to 2D, highlighting the need for more accurate tissue-mimetic TM model systems for drug screening purposes (Bhadriraju and Chen, 2002; Edmondson et al., 2014). Our recently described HTM cell-encapsulated ECM hydrogel system was engineered with this in mind (Li et al., 2021). The HTM hydrogel facilitates accurate and reliable modeling of 3D cell-cell and cell-ECM interactions as seen in the juxtacanalicular region of the native TM tissue, at a level not possible with other biomaterials-based models (for a recent review see (Lamont et al., 2021)). Importantly, this can be accomplished under both normal and simulated glaucomatous conditions (Li et al., 2021; Li et al., 2022a; Li et al., 2022b). We previously demonstrated the bidirectional responsiveness of the HTM hydrogel to pathologic TGFβ2 induction (Inatani et al., 2001) and ROCKi rescue using standard Y27632 (Rao et al., 2001) for correlative analyses of TM cell cytoskeletal organization with tissue-level functional changes (Li et al., 2021).

Therefore, the goal of the present study was to use our HTM hydrogel as screening platform to investigate the effects of clinically-used netarsudil and different netarsudil-family experimental ROCKi compounds on reversing TGFβ2-induced pathologic HTM cell contraction and actin stress formation – both strongly associated with outflow dysfunction in glaucoma. Unlike other ocular hypertension/glaucoma medications that do not specifically target the diseased outflow tract, netarsudil directly addresses the TM outflow pathway. Its main mode of action is to increase outflow via disassembly of TM cell ECM-focal adhesions as well as actin stress fibers, thereby potently relaxing the stiffened tissue (Zhang et al., 2012; Wang and Chang, 2014; Sturdivant et al., 2016; Rao et al., 2017; Lin et al., 2018; Tanna and Johnson, 2018). In addition to the clinically-used netarsudil, two sets of experimental ROCKi compounds related to netarsudil were selected and provided by Aerie Pharmaceuticals Inc. One set included two compounds that performed similarly or better than netarsudil in Aerie’s *in vitro* screening but did not perform as well in intraocular pressure-lowering studies using normotensive Dutch Belted rabbits (unpublished data): compounds A and C. The other set included two compounds that did not perform as well as netarsudil both *in vitro* and *in vivo*: compound B and D.

First, we investigated the specific inhibitory activity of the different netarsudil-family ROCKi compounds against ROCK1 and ROCK2 together with their ability to disrupt HTM cell focal adhesions. Of note, these biochemical and cell-based activity assays were performed independently in the laboratories of Aerie Pharmaceuticals. All ROCKi compounds exhibited significant ROCK1/2 inhibitory activity, with the order of potency (K_i (average of ROCK1/2)_) from highest to lowest being: compound A (~1.4 nM) >compound C (~1.5 nM)>compound B (~1.9 nM)>netarsudil (~2.7 nM)>compound D (~3.1 nM). Similarly, the rank order of potency (IC_50_) to disrupt HTM cell focal adhesions was: compound C (~5.6 nM)>compound A (~15 nM)>netarsudil (~17 nM)>compound D (~193 nM). Compound B was not directly tested; however, in a previous experiment its racemate (i.e., 50% of compound B and 50% of its inactive enantiomer) was assessed in the HTM cell focal adhesion assay showing an IC_50_ of ~42 nM (unpublished data). With this information in mind, the expanded rank order would be: compound C>compound A >netarsudil>*compound B*>compound D – clearly grouping compounds A and C in a similar range as netarsudil, whereas compounds B and D overall would group below netarsudil. Collectively, all netarsudil-family ROCKi were far more potent than Y27632 (K_i_ of ~31.5 nM; IC_50_ of ~1738 nM), consistent with previous studies using the same biochemical and cell-based assays (Lin et al., 2018). The differential responses observed with clinically-used netarsudil compared to experimental compounds A-D likely stems from differences in compound structure/chemistry, affecting cellular uptake and efficacy. Compound A was nearly identical to netarsudil in terms of disrupting HTM cell focal adhesions but as its benzoic acid moiety is unsubstituted, it is more quickly metabolized than netarsudil. The metabolism occurs when its ester linkage is hydrolyzed by cellular esterases. In contrast, compounds B and C cannot be metabolized by cellular esterases at all. And while compound D potently inhibited ROCK1/2 activity, it is partially ionized at physiological pH and performed worst in the HTM cell focal adhesion assay. It is thought that this ionization inhibits cellular penetration and thus decreases the compound’s effectiveness in whole-cell assays.

Next, we investigated the dose response behavior of the different netarsudil-family ROCKi compounds in rescuing pathologic TGFβ2-induced HTM hydrogel contraction according to our previously published protocol (Li et al., 2021). All ROCKi compounds exhibited potent contraction-reversing effects on HTM cell-encapsulated ECM hydrogels upon glaucomatous induction in a dose-dependent manner. The order of potency from highest to lowest was: compound A (EC_50_ of 3.7 nM)>compound C (6.7 nM)>compound B (22.2 nM)>netarsudil (35.9 nM)>compound D (123.3 nM). Strikingly, this precisely matched the compounds’ independently determined ROCK1/2 inhibitory activity profiles, which were made available to the masked study team only after the completion of all hydrogel experiments to avoid unintentional bias. Importantly, these data suggest that the HTM hydrogel contraction assay based on a robust glaucomatous induction and rescue protocol facilitates accurate measurements of ROCKi potency in a tissue-mimetic 3D ECM microenvironment. The subtle differences observed in the rank order of netarsudil-family ROCKi potency between the HTM hydrogel contraction assay (for EC_50_) and the HTM cell focal adhesion disruption assay (for IC_50_) could be explained by the differences in cell source. While all hydrogel studies herein used primary HTM cells derived from healthy donors, the focal adhesion assay relied on the SV40 TAg transformed HTM cell line TM-1 (Filla et al., 2002; Liu et al., 2002).

We then directly compared the different netarsudil-family ROCKi compounds at their respective EC_50_ to assess the compounds’ tailored efficacy in rescuing TGFβ2-induced HTM hydrogel contraction and actin stress fiber formation within the 3D biopolymer network. First, the different ROCKi treatments exhibited distinct signatures of reversing pathologic HTM hydrogel contraction independent of the cell strain used. The overall order of efficacy from highest to lowest was: netarsudil (relative hydrogel size vs. controls of ~120%; TGFβ2 at ~75%)>compound B (~105%)>compound D (~96%)>compound C (~87%)>compound A (~78%). These findings were somewhat unexpected as they did not match the rank order of the earlier dose response experiments. The same ROCKi stock solutions were used to prepare the distinct EC_50_-level treatments as single-use aliquots to avoid potential freeze and thaw damage. Hence, there were no concerns of drug degradation. However, it is possible that the aggressive two-step dilution for compounds A and C, which performed worst in this experiment, from the original 1 mM stock to obtain the desired low nanomolar doses may have contributed to drug instability in the organic solvent DMSO. All other ROCKi compounds required only a one-step dilution to achieve the EC_50_-level doses, lending support for this potential explanation. Of note, these data are in line with Aerie’s observation of compounds A and C being unable to significantly lower intraocular pressure in studies using normotensive Dutch Belted rabbits (unpublished data). A key take-away from these experiments is that netarsudil at both EC_50_ and maximum efficacy level outperformed the experimental ROCKi compounds A-D, thereby validating its clinical status in our 3D TM hydrogel system. Second, the different netarsudil-family ROCKi compounds exhibited distinct effects of relaxing pathologic F-actin stress fibers in ECM hydrogel-encapsulated HTM cells. The order of overall efficacy from highest to lowest was: compound D (relative F-actin signal fold-change vs. controls of ~0.8; TGFβ2 at ~1.9)>netarsudil (~1.0)>compound B (~1.1)>compound C (~1.3)>compound A (~1.7).

Hence, the actin cytoskeletal remodeling data also revealed a complex pattern in which the most potent compounds A and C according to their EC_50_ were the worst performers here. Collectively, these data showed that netarsudil and most/all other related experimental ROCKi compounds potently rescued pathologic HTM hydrogel contraction and relaxed fibrotic-like F-actin stress fibers, consistent with previous studies (Lin et al., 2018; Keller and Kopczynski, 2020).

Lastly, to eliminate the potential confounder of variable ROCKi dosing, we compared the efficacy of the different netarsudil-family compounds in rescuing TGFβ2-induced HTM hydrogel contraction at uniform netarsudil-EC_50_-levels across treatment groups. To do so, the dosing of the various compounds was adjusted as follows: compound A (~10-fold more), compound B (~1.6-fold more), compound C (~5.3-fold more), and compound D (~3.4-fold less). When tested at equimolar dosing, we observed that netarsudil performed similarly to the related experimental ROCKi compounds A-D independent of the HTM cell strain used, suggesting that subtle differences in compound-specific activities can be offset by matching their doses with the clinical drug.

In conclusion, our data suggest that (i) netarsudil exhibits high potency to reverse pathologic TM cell contraction and actin stress formation in a tissue-mimetic 3D ECM microenvironment in support of its clinical use, and (ii) that our bioengineered hydrogel is a viable screening platform to complement and expand conventional 2D monolayer cell cultures and preclinical animal models in pursuit of furthering our understanding of TM cell pathobiology in glaucoma.

## Data and materials availability

All data needed to evaluate the conclusions in the paper are present in the paper and/or the Supplementary Materials. Additional data related to this paper may be requested from the authors.

## Acknowledgments

We thank Dr. Robert W. Weisenthal and the team at Specialty Surgery Center of Central New York for providing corneal rim specimens. We also thank Dr. Nasim Annabi at the University of California – Los Angeles for providing the KCTS-ELP, and Dr. Mariano S. Viapiano and the Neuroscience Microscopy Core at Upstate Medical University for imaging support.

## Author contributions

T.B., A.S., R.G., H.Y., C.K., M.A.d.L., C.C.K., and S.H. designed all experiments, collected, analyzed, and interpreted the data. T.B. and S.H. wrote the manuscript. All authors commented on and approved the final manuscript. S.H. conceived and supervised the research.

