## Supplemental information for "Effects of netarsudil-family Rho kinase inhibitors on human trabecular meshwork cell contractility and actin remodeling using a bioengineered ECM hydrogel"

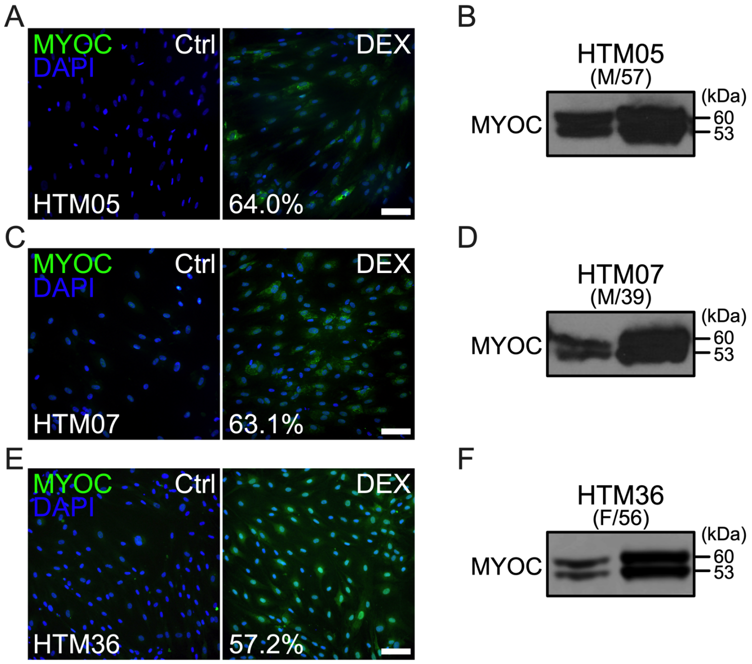


**Suppl. Fig. 1. HTM cell characterization.** (**A**) Representative fluorescence micrographs of intracellular myocilin (MYOC) at 7 d with percent dexamethasone (DEX)-induction. Scale bar, 100 µm. (B) Immunoblots of secreted MYOC at 7 d.

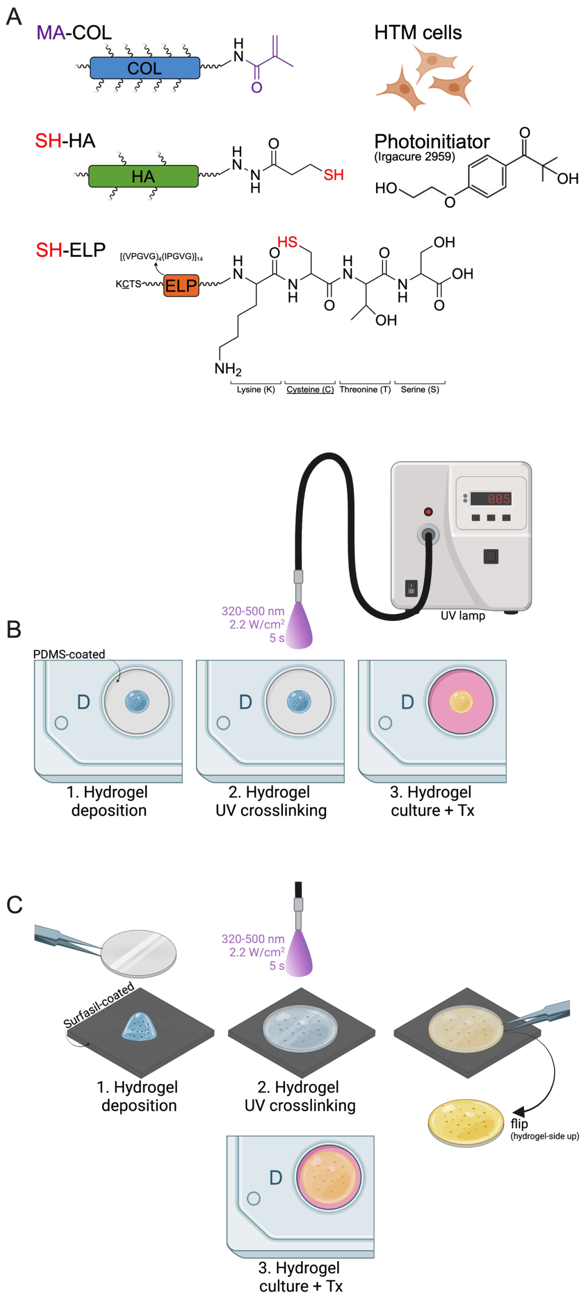


**Suppl. Fig. 2. Schematic of ECM biopolymer precursors and hydrogel formation.** (**A**) Schematic of methacrylate-conjugated collagen type I (MA-COL), thiol-conjugated elastin-like polypeptide (SH-ELP; elastic ($[VPGVG]_4[IPGVG])_{14}$ core sequence, thiol via K<u>C</u>TS flanks), and thiol-conjugated hyaluronic acid (SH-HA). HTM hydrogels were fabricated by mixing HTM cells ($1x10^6$ cells/ml) MA-COL, SH-HA with photoinitiator, and in-house expressed SH-ELP. (**B**) 10 µl of the HTM cell-containing hydrogel precursor solution were plated on PDMS-coated 24-well plates, or (**C**) 30 µl were plated on Surfasil-coated 18x18-mm square coverslips followed by placing regular 12-mm round coverslips atop. All HTM hydrogels were UV crosslinked (320-500 nm, 2.2 W/cm$^2$, 5 s) and cultured in growth media in presence of the different treatments for 10 d. Created with BioRender.com.

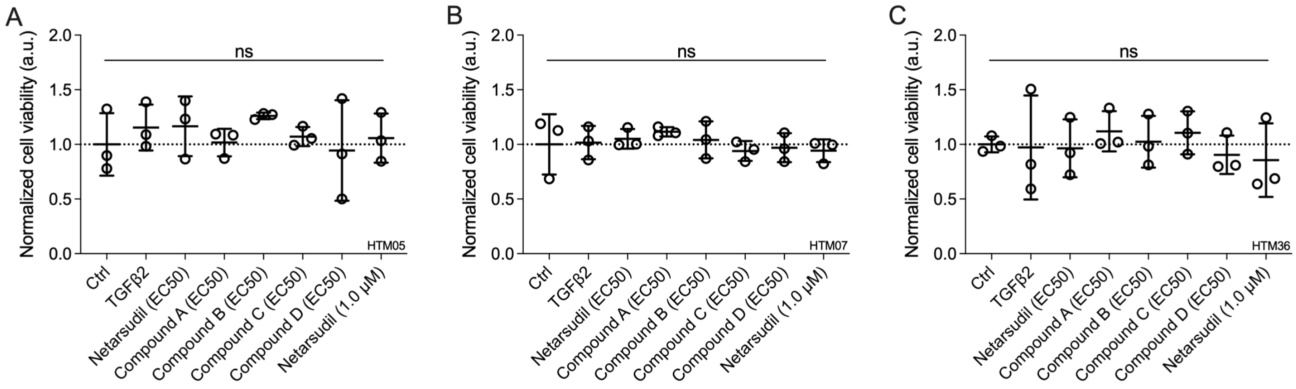


**Suppl. Fig. 3. Effects of netarsudil-family ROCKi treatment following TGFβ2-induction on HTM cell viability within ECM hydrogels.** Cell viability quantification of HTM hydrogels encapsulated with (**A**) HTM05, (**B**) HTM07, or (**C**) HTM36 subjected to the different treatments for 10 d (N = 3 replicates per group and HTM cell strain). Data shown as Mean ± SD with individual data points. Significance was determined by one-way ANOVA using multiple comparisons tests (ns = not significant).

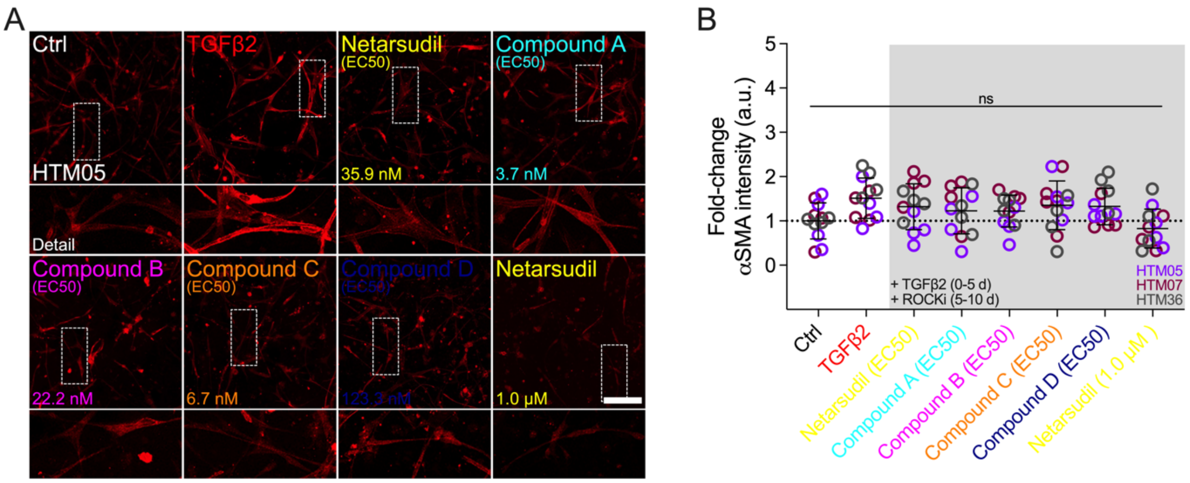


**Suppl. Fig. 4. Effects of netarsudil-family ROCKi treatment following TGFβ2-induction on HTM cell alpha smooth muscle actin (αSMA) fibers within ECM hydrogels.** (**A**) Representative confocal fluorescence micrographs of αSMA in HTM hydrogels encapsulated with HTM05 subjected to the different treatments for 10 d. Scale bar, 200 μm. (**B**) Pooled quantification of relative αSMA signal intensity in HTM hydrogels encapsulated with HTM05 (purple), HTM07 (maroon), or HTM36 (gray) (N = 4 replicates per group and HTM cell strain). Data shown as Mean ± SD with individual data points. Significance was determined by two-way ANOVA using multiple comparisons tests (ns = not significant).
